# Elucidating the Mechanical Energy for Cyclization of a DNA Origami Tile

**DOI:** 10.1101/2021.02.07.430115

**Authors:** Ruixin Li, Haorong Chen, Hyeongwoon Lee, Jong Hyun Choi

## Abstract

DNA origami has emerged as a versatile method to synthesize nanostructures with high precision. This bottom-up self-assembly approach can produce not only complex static architectures, but also dynamic reconfigurable structures with tunable properties. While DNA origami has been explored increasingly for diverse applications such as biomedical and biophysical tools, related mechanics are also under active investigation. Here we studied the structural properties of DNA origami and investigated the energy needed to deform the DNA structures. We used a single-layer rectangular DNA origami tile as a model system and studied its cyclization process. This origami tile was designed with an inherent twist by placing crossovers every 16 base-pairs (bp), corresponding to a helical pitch of 10.67 bp/turn which is slightly different from that of native B-form DNA (10.5 bp/turn). We used molecular dynamics (MD) simulations based on a coarse-grained model on an open-source computational platform, oxDNA. We calculated the energies needed to overcome the initial curvature and induce mechanical deformation by applying linear spring forces. We found that the initial curvature may be overcome gradually during cyclization and a total of ~33.1 kcal/mol is required to complete the deformation. These results provide insights into the DNA origami mechanics and should be useful for diverse applications such as adaptive reconfiguration and energy absorption.

## INTRODUCTION

DNA self-assembly is as a powerful bottom-up manufacturing approach based on spontaneous sequence complementarity.^1,2^ DNA nanotechnology can produce complex architectures with subnanometer accuracy and precision, using DNA tiles^3–5^ and DNA bricks^6–8^. The tile-based assembly uses several short single-stranded DNA (ssDNA) oligonucleotides to assemble one or few motifs, which then form two-dimensional (2D) or three-dimensional (3D) configurations by using their sticky ends. In contrast, each DNA brick is a unique ssDNA strand, thus more intricate geometries may be constructed from a single step annealing reaction. Dynamic DNA nanostructures are also available, including tweezers^9,10^ and walkers^11–14^. These mobile nanodevices rely on toehold-mediate strand displacement reaction,^15–17^ to detach from some of the strands in order to stay undefined or bind with new strands. The major difference between the two devices is the designed movement; DNA tweezers mostly switch between open and closed states, while DNA walkers can move along a track as long as fuel strands are available.

DNA origami is a widely used method due to excellent programmability and structural predictability.^18–23^ In this approach, a long ssDNA ‘scaffold’ strand (up to several thousand basepairs or bp) is laid out in a folded pattern to approximate the target design. A number of short ssDNA ‘staple’ strands are then added to bring selected parts of scaffold together by hybridization to form designed conformations. Various 2D and 3D DNA origami structures have been demonstrated, including shafts,^24–26^ tiles,^27–30^ cubes,^31–33^ and wireframe polygons^34–36^, to name a few. Dynamic, shape-changing DNA origami structures were also realized via two-step reactions: strand displacement and reannealing.^37–39^ While the strand displacement reaction is possible at room-temperature due to the low energy barrier, the reannealing with new strands is often performed at elevated temperatures to provide necessary activation energies.^40–42^ For both static and dynamic DNA origami architectures, their structural properties and mechanical behaviors are of great interest, including the twisting of a double-stranded DNA (dsDNA) bundle,^43,44^ the interconnection between bundles by crossovers,^45^ the interaction between ds-regions in an origami structure,^46^ and the structural deformation by applied external forces.^47^ A better understanding of such fundamental mechanics will significantly advance the design and development of DNA origami for a variety of applications.

There are two typical lattices (honeycomb and square) for cross-section designs of DNA origami. These two design approaches may be distinguished by crossover arrangements as they are distinct and different based on the layouts of dsDNA bundles and crossovers. In a honeycomb lattice, a crossover is placed every 2/3 turn. Given the helicity of 10.5 bp/turn for native B-form DNA, it is exactly 7 bp per crossover.^48,49^ In contrast, a crossover is placed every 3/2 turns in a square lattice, yielding 15.75 bp per crossover. However, only an integer number of base pairs is possible in the design. Thus, 16 bp per crossover is used frequently, which corresponds to 10.67 bp/turn. The difference between 10.67 and 10.5 bp/turn in DNA helicity causes a right-handed twist in the entire structure.^20,24,27^ The structural twist and curvature may be resolved by forcing the upper and lower boundaries together using linker staples^27^ or by adding intercalators that directly modulate the helicity (*e.g*., from 10.5 to 10.67 bp/turn).^24,50–52^ Regardless of the methods, external (mechanical or chemical) energy will be accounted for the structural correction, and this is the activation energy required for the deformation of the origami.

In this work, we investigated a cyclization process of a single-layer DNA origami rectangle. This origami tile is designed in a square lattice and thus has an inherent structural curvature. In a previous publication,^27^ we used this tile in a series of experiments to study the energy needed for cyclization as functions of incubation temperature, origami size, and structural defect. The atomic force microscopy (AFM) measurements and mechanical analysis of elastic deformation were performed to provide insights into the reconfiguration, including relevant reaction rates and the bending spring constant of crossovers. While the experiment and elasticity theory revealed the overall aspects of the origami deformation, the detailed process was not known, because only the DNA conformations before and after the cyclization reaction were available. In the current study, we performed numerical simulations of the DNA origami tile cyclization by directly applying external forces. The multistep simulations of small, quasi-equilibrium deformations allowed us to elucidate the details of the entire cyclization process and calculate the mechanical energy associated with conformational change at each step. The computation was performed with coarsegrained molecular dynamics (MD) model in oxDNA, an open-source computational platform. The MD computation was also complemented by finite element method (FEM) simulations in CanDo, a free, online simulation software. The evolution of the simulated structure in each state shows how the DNA origami tile transforms gradually from its initial twisted conformation to the final cyclized configuration. The overall energy for the complete cyclization can be calculated by summing all the mechanical energies in the multistep simulations and is approximately 33.1 kcal/mol. In addition, we also calculated the energy needed to flatten the twisted DNA origami tile, which is ~26.7 kcal/mol. The energies for cyclization and flattening are in excellent agreement with previous experimental and theoretical results.^27^

## MATERIALS AND METHODS

### Origami Tile Design

Figure 1 shows the design of a single-layer DNA origami rectangle used as a model system in this computational study. There are 32 helices in height (*i.e*., 32 rows of dsDNA bundles) and 224 bases in width (in each row). The origami tile is sketched in caDNAno2^48^. To avoid blunt-ended stacking, only 192 bases in width are used and the 16 nucleotides (nt) on either side are left unpaired. The sequence information is available in our previous report.^27^ The designed width is calculated to be approximately 64 nm based on 0.332 nm/bp along the axis of regular, right-handed B-DNA. The height is estimated using the diameter of a dsDNA bundle (~2.2 nm) and the distance between the center of neighboring bundles (~2.3 nm when closely packed).^20,46^ However, the gap between adjacent bundles is noticeably wider than the bundle diameter based on our previous AFM results.^27^ For a fair estimation, we assume the gap to be approximately 2.3×1.2 nm. Thus, the designed height is estimated as ~88 nm. A set of linker staples, if introduced in an experiment, can hybridize with upper and lower boundaries (rows 1 and 32) simultaneously, thereby cyclizing the tile into a tube, as we demonstrated in a previous report.^27^ Experimentally, the DNA tile experiences thermal fluctuation that deforms the structure randomly, and the linkers seal the upper and lower edges when they are in proximity. By design, the cyclized tube is approximately 64 nm in width and 28 nm in diameter. If collapsed to a flat surface, the origami will be about 64 nm in width, 44 nm in height and 4 nm in thickness (double layer). The AFM results suggest that the synthesized tile measures about 90×65×2 nm and the flattened tube is around 45×65×4 nm.^27^

**Figure 1.**
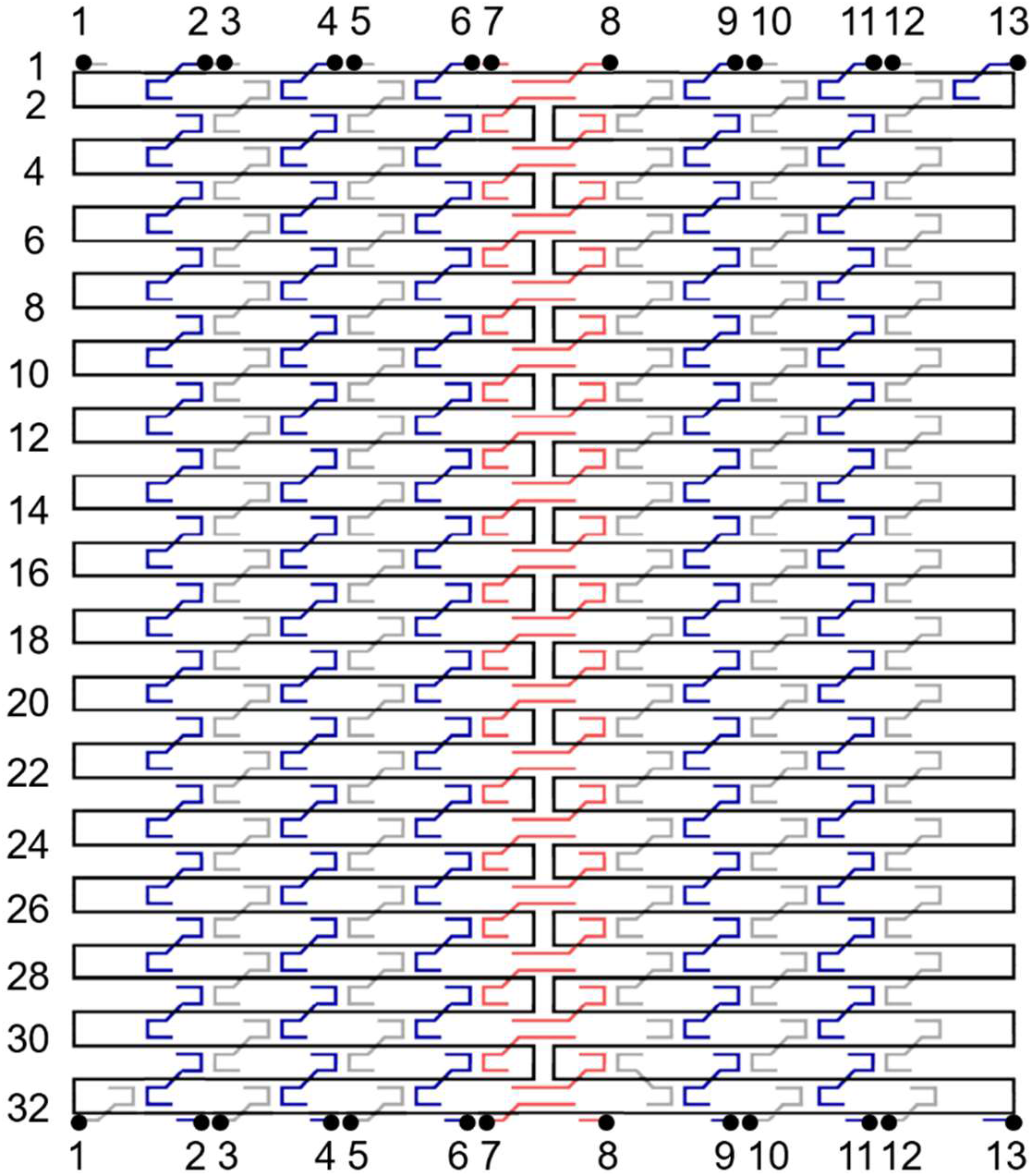
Folding path diagram of a DNA origami tile. Black line denotes the scaffold strand, which routes clockwise (5’ to 3’ end). Blue, gray, and red lines represent staples. The color marks different shape and directions. Blue and gray staples have the same shape but opposite directions: blue goes down while gray goes up from 5’ to 3’ end. The red staples secure the scaffold in the middle of the origami and thus have a different shape compared with blue and gray. A typical staple length is 32 nt. The dsDNA bundles are numbered 1 through 32, as noted on the left. Note that a total of 13 gray and blue staples were used to seal the upper and lower edges to form a cyclized tube in the experiment. The 13 pairs of the black dots indicate the cut sites of the linker staples. This design was used in the MD simulations.

The DNA origami tile is designed in a square lattice, with a crossover every 16 nt. Typically a staple is 32 nt in length – the middle 16 nt binds with a dsDNA helix while the 8 nt parts at both 5′ and 3′ ends bind with neighboring two dsDNA bundles. This is equivalent to a helicity of 10.67 bp/turn. Therefore, the tile will have a slight, right-handed global twist if subjected under 10.5 bp/turn conditions. In the simulations, 13 linker staples designed to cyclize the DNA origami rectangle were cut into halves as illustrated in Figure 1. The cutting sites are marked with 13 pairs of black dots and labelled 1 through 13. The origami tile can then be cyclized by applied external forces.

### FEM Simulations with CanDo

FEM is a numerical method for solving partial differential equations (PDEs) in 2D or 3D space variables. The method subdivides a large system into smaller, simpler parts that are called finite elements to solve the equations. In our case, FEM simulations were performed with CanDo on single base-pair resolution to study 3D equilibrium conformations of the DNA origami tile (without linker staples). The axial rise per base-pair and the helix diameter were set at 0.34 nm and 2.25 nm. The crossover spacing was 10.5 bp. All the mechanical properties of dsDNA were set as default: axial stiffness at 1100 pN, bending stiffness at 230 pN·nm^2^, torsional stiffness at 460 pN·nm^2^. To simplify the properties of ssDNA, nick stiffness factor was set at 0.01, which accounts for the property ratio between ssDNA and dsDNA. For example, the axial stiffness of ssDNA is 0.01×1100 pN, which is 11 pN. The simulations yield an inherent global twist in the origami conformation.

### MD Simulations with OxDNA

MD computation is a method for analyzing the physical movements of atoms and/or molecules (particles). All-atom models include all the related atoms in the simulations for better accuracy, while coarse-grained models use a pseudo-atom to represent a group of atoms to save simulation time. The particles are subject under interactions for a fixed duration of time, giving a view of the dynamic evolution of the system. The trajectories of the particles are typically determined by numerically solving Newton’s equations of motion for the particles.

The MD simulations based on a coarse-grained model were performed with oxDNA^53^ to study equilibrium conformations of the DNA origami tile and their gradual structural deformations in series during cyclization and flattening by external loading. Specifically, the oxDNA2 model was used with the salt concentration set at [Na^+^] = 0.5 M and the temperature at 300 K.^54^ The caDNAno2 file of the origami tile was converted into topology and configuration files as the initial conformation in oxDNA with actual sequence information. To simplify the computation, all the ss-regions that do not serve any structural purposes (e.g., 8-nt toeholds on linker staples and free scaffold loops) were not included in the initial conformation. Since these regions were barely seen interfering with dsDNA bundles under AFM, it is reasonable to exclude them. As discussed in the ‘Origami Tile Design’ section, the linker staples were cut into two halves so that the upper and lower boundaries were separate before structural deformation. The configuration was relaxed by MD simulations without any external forces for 2×10^6^ steps to obtain the initial equilibrium structure.

In the cyclization process, external linear spring forces were introduced by mutual traps in oxDNA (see oxDNA documents for details) and applied on the cut sites of the linkers (13 pairs of black dots located at the upper and lower boundaries in Figure 1) on origami tile to induce a cyclization through a series of deformations. The potential energy associated with the spring forces was determined by considering the conformation in each state. Figure 2a shows the loading on a pair of nucleotides. A hypothetical spring that connects nucleotide A and B has a spring force constant *k* and an equilibrium length *l*_0_. Given the distance of A and B is *l*, the forces can be expressed following the Hooke’s law.

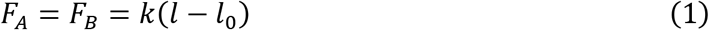

**Figure 2.**
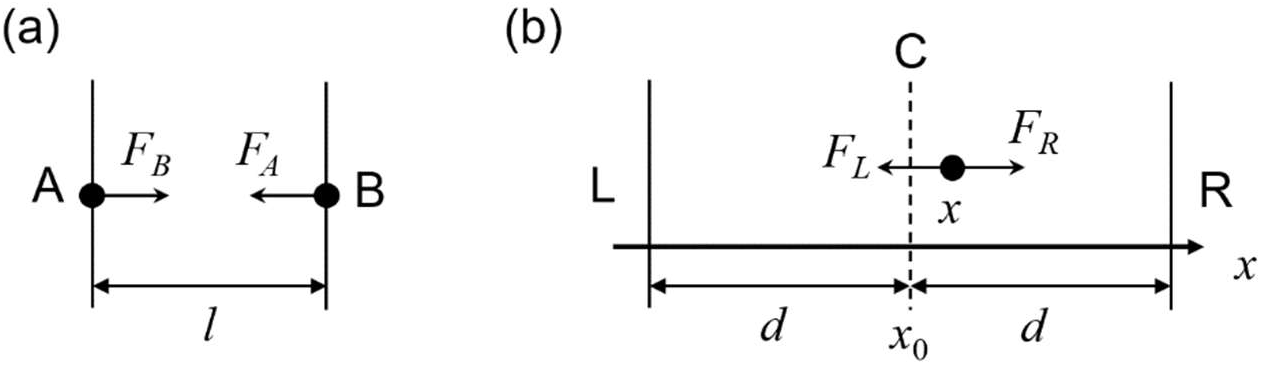
Mechanical loadings in oxDNA simulations. (a) Hypothetical linear spring forces between two nucleotide A and B (solid dots) with a distance *l*. The forces on A pointing to B is denoted as force *F_B_*, and similar naming applies to *F_A_*. A set of spring forces was used for cyclization of the DNA origami tile. (b) Hypothetical linear spring forces pulling a nucleotide (solid dot) at *x* to an imaginary plane (dashed line, marked as C) located at *x*_0_. The forces are realized by two planes, L and R (vertical lines), in a distance *d* from the imaginary plane on the left and right sides. The force toward the left plane (L) is *F_L_*, whereas the force toward the right plane (R) is *F_R_*.

Assuming the potential energy is 0 at *l* = *l*_0_, the potential energy of the spring system can be calculated.

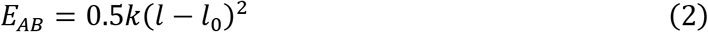

For a given conformation of the origami tile, *l* is set. The potential energy can be modulated by varying *k* and *l*_0_ (i.e., changing the spring) in each calculation from a state to the next. Two million calculation steps were carried out, which took about two days of computation. After each calculation, the results were analyzed by visualizing the structures in cogli2 and by extracting the directional and positional data. The conformation with the minimum of global potential energy was chosen as the next state. The distance between two nucleotides from the cut of each linker staple was measured from the positional data to obtain a vector pointing from one terminal to the other.

In the flattening process, external linear spring forces were introduced by two force planes in oxDNA (see oxDNA documents for details) and applied on all 13,360 nucleotides of the origami tile to force them to an imaginary center plane through a series of deformations. If the plane is aligned perpendicular to the x-axis, the loading of any single nucleotide will be as shown in Figure 2b. We placed two force planes for loading. The force toward the left plane (L) is *F_L_*, whereas the force toward the right plane (R) is *F_R_*. Each plane exerts a spring force constant k in a distance *d* from the center plane (dashed line, C). Given that the positions of the center plane and the nucleotide are *x*_0_ and *x*, respectively, the forces are

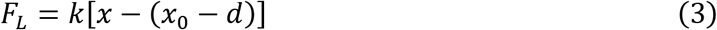

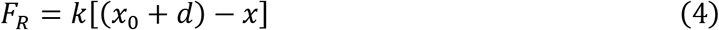

Note that the directions of the forces are marked in Figure 2b. If the positive direction is set as the direction of the *x*-axis, the total force is

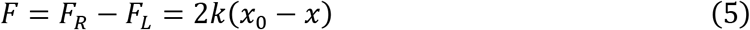

Thus, the actual direction of the total force *F* is always pointing to the center plane and the magnitude is proportional to the distance of the nucleotide from the center plane. Note that the force is independent of *d*. Similar to the two nucleotides linked by a spring force pair as shown in Figure 2a, the potential energy of the spring system may be determined with an assumption that the potential energy is 0 at *x* = *x*_0_.

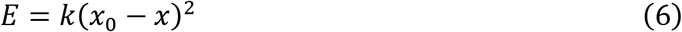

For a given origami conformation, *x* is set. The location of the center plane is determined by the initial state (state 0), which means that *x*_0_ is set as well. The potential energy will then be modulated only by assigning different *k* value (*i.e*., different spring). The potential energy of the forces, the states, and the distance between each nucleotide and the imaginary plane were determined in a similar manner as in the cyclization process.

## RESULTS

### Experiments and Theory

Figure 3a and b illustrates two, honeycomb and square, lattices in typical DNA origami designs. In a honeycomb lattice, a crossover is placed every 2/3 turn, which corresponds to exactly 7 bp per crossover. This crossover arrangement is compatible with the intrinsic helicity (10.5 bp/turn) of right-handed B-form dsDNA. In contrast, a crossover is placed every 3/2 turns in a square lattice, which equals to 15.75 bp per crossover. The closest integer 16 bp per crossover is typically used, thus yielding 10.67 bp/turn. This will cause a right-handed twist in the global structure. Figure 3c shows the conformation of our DNA origami tile from the FEM simulations with 10.5 bp/turn. A slight, right-handed curvature is observed, thus matching with the lattice design. If the tile is cyclized and the upper and lower edges are sealed with a set of linker staples (shown as the purple line), the structure will have all the dsDNA bundles (shown as gray rods) perfectly aligned as illustrated in Figure 3d.

**Figure 3.**
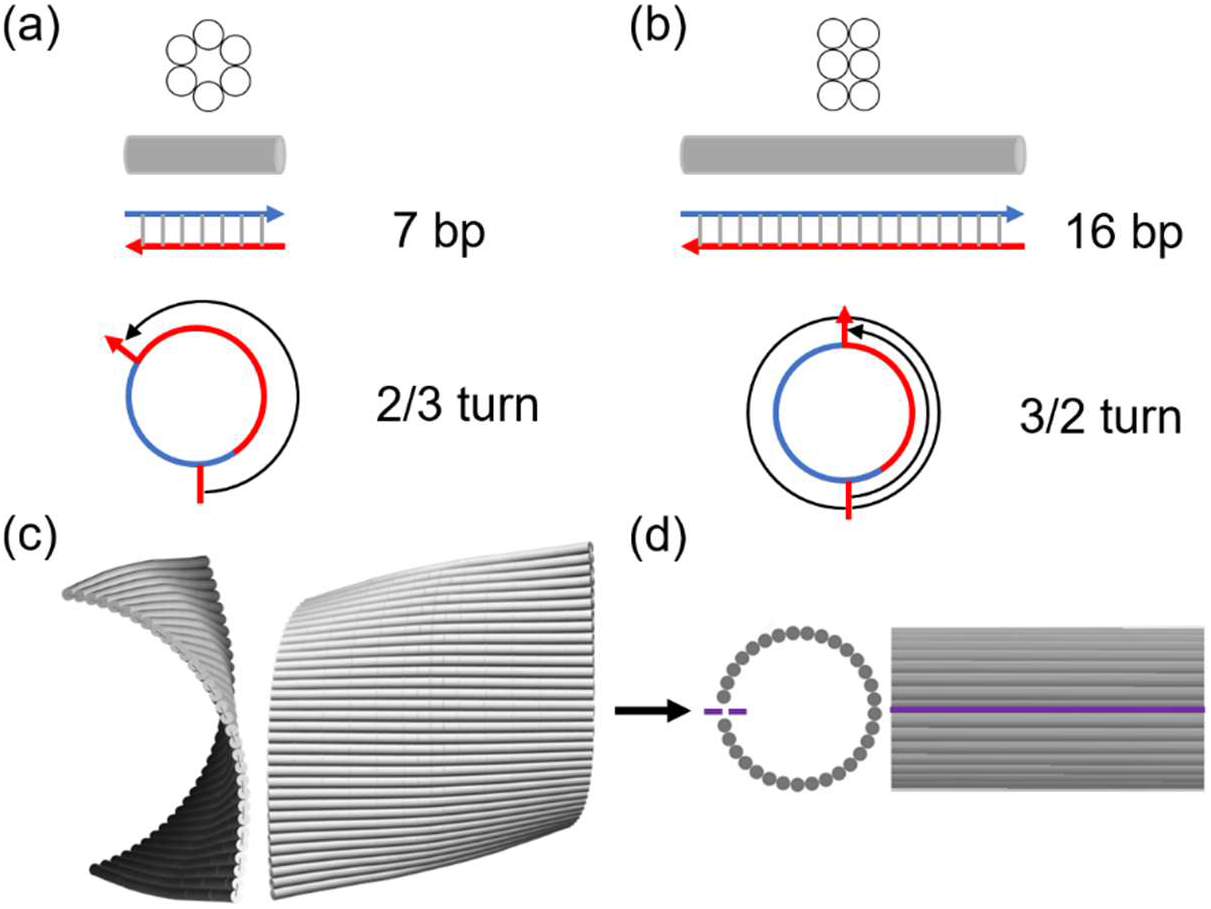
Schematics and FEM simulation results of DNA origami. Comparison of (a) honeycomb and (b) square lattice designs. The gray cylinder represents a dsDNA bundle from the lattice. The blue and red lines with arrows indicate the scaffold and staple strands, respectively, with their 3’ ends marked as arrows. In a honeycomb lattice, crossovers can be placed every 7 bp which is accounted for 2/3 turns in the axial direction of a dsDNA bundle. This corresponds to 10.5 bp/turn, consistent with the helicity of native B-form DNA. In a square lattice, however, a possible crossover is arranged in every 16 bp or 3/2 turns, yielding 10.67 bp/turn. (c) CanDo simulation results of a single-layer DNA origami rectangle in a square lattice (side and front views). With the designed helicity at 10.67 bp/turn, the tile has a minor global twist in the FEM simulations at 10.5 bp/turn. (d) Schematics of the cyclized DNA origami tile in both views. The purple line indicates the sealed upper and lower boundaries by linker staples in experiments.

It is worth noting that the curvature of the DNA origami tile before cyclization cannot be observed under AFM. During deposition, the DNA origami collapses on the mica surface by the attraction force (in the presence of metal cations such as Mg^2+^), thus flattening the tile.^27^ To see the curvature clearly, multiple tiles may be polymerized side by side into a long ribbon (> 1 μm in the polymerized direction). We previously found that the ribbon also flattens under AFM, but the accumulated twisting leads to a multiple folding of the ribbon and parallelogram-shaped kinks emerge at the folded parts.^24^ Moreover, the kinks indicate the handedness of the curvature and their density may change if the DNA helicity is altered. For example, the right-handed twist can be weakened, flattened, and even overturned to a left-handed conformation by modulating from 10.5 to 10.67 and then to 10.84 bp/turn.

We studied the cyclization of the single-layer DNA origami rectangle designed at 10.67 bp/turn in a series of experiments as well as theoretical calculations in a previous publication.^27^ In the experiments, we added 13 linker staples to the solution containing preformed DNA origami tiles and held at various temperatures from 35 °C to 50 °C. Then, we measured the origami conformations after different durations of incubation, such as 0.5, 1, 2, and 4 hours. The fractions were determined by counting the origami species based on their conformations (*e.g*., flat vs. cyclized tiles) in the AFM images. A set of ordinary differential equations (ODEs) was developed to relate the fractions of species as functions of incubation time and temperature. Several kinetic parameters such as reaction rate constants were determined by numerically fitting the experimental data. Along with rate constants, the activation energy of 32.4 ± 0.7 kcal/mol was obtained based on the Arrhenius plot.^27^ This is the minimum energy needed to overcome barriers and drive the cyclization.

The cyclization process was also studied theoretically.^27^ Due to the difficulties in the theoretical calculation of cyclization, the structural transformation was conceptually analyzed as a two-step (flattening and flexing) process. First, the global twist was resolved hypothetically to form a perfectly planar structure by forcing the helicity from 10.5 to 10.67 bp/turn. The twist angle of 8.23° per 16 bp was calculated considering the difference in the designed and natural DNA helicity (10.67 and 10.5 bp/turn). Given spring constants of dsDNA bundles, the energy needed for flattening the tile was calculated as ~25.6 kcal/mol. Next, the planar structure was bent into an ideally circular shape (as shown in the Fig. 3d). This was realized by twisting of dsDNA bundles and bending of crossovers. These two components were modeled as elastic springs connected into a network whose effective spring constant was calculated by considering the springs in parallel and series. With knowledge of the effective spring constant of the entire origami, the energy necessary for bending at 360° was approximately 6.7 kcal/mol. Combining the two steps, the origami tile transformed from a twist conformation to a planar tile and then to a cyclized tube. Thus, the driving energy for the cyclization process was acquired by summation of the energies for flattening and flexing, yielding ~32.3 kcal/mol.^27^ This theoretical value agrees almost perfectly with the experimentally determined activation energy.

### Mechanical Energy Induced Cyclization

Coarse-grained MD simulations were performed to analyze the detailed cyclization process. We realized a series of quasi-equilibrium deformations by defining multiple states and calculating potential energies from one state to the next. First, an equilibrium structure was set as the initial state (state 0). Later states were named from 1 to 10 as shown in Figure 4a. The structure was subject under hypothetical linear spring forces in each process from one state to the next. From state 0 to state 1, the forces were added between neighboring dsDNA bundles to create the starting conformation of the cyclization (state 1, see Figure S1). This step was needed since the tile may fold in a random direction toward either side upon external forces, thus the cyclization may not be achieved. For state 1 to 2, 2 to 3, and all the rest until the final state, the forces were added between 13 pairs of cut sites on the linker staples (shown as dots in Figure 1). By pulling the linkers, the upper and lower edges will be brought together, cyclizing the origami tile.

**Figure 4.**
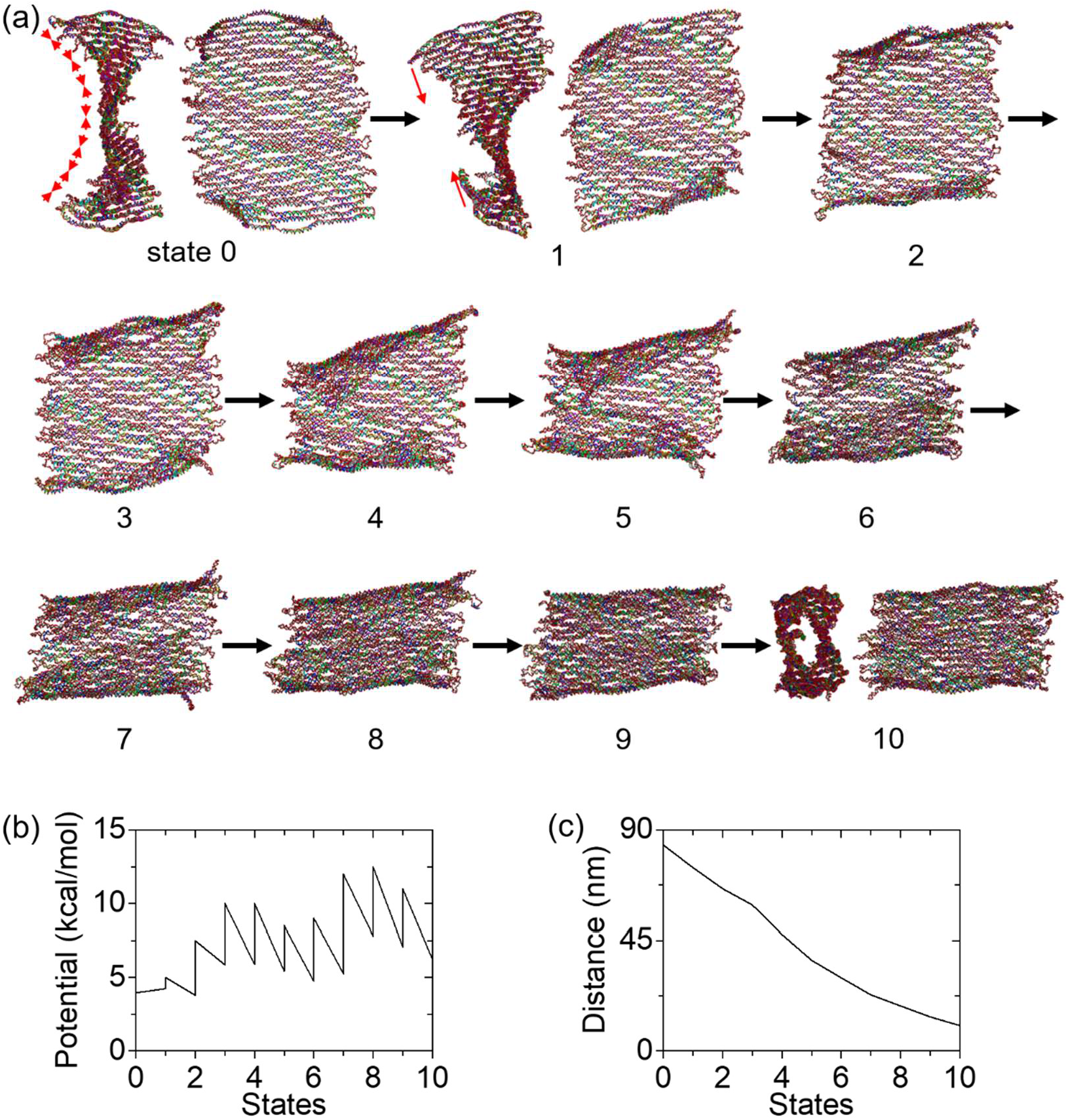
Deformation process of origami tile cyclization in MD simulations. (a) The DNA tile deforms from state 0 (initial) to state 10 (final). In the final state, the tile is cyclized into a tube. The cyclization is induced by adding hypothetical linear spring forces in the MD simulations. From state 0 to state 1, the forces were added between neighboring dsDNA bundles (illustrated by the red arrows in state 0) to create the starting conformation for the cyclization (state 1). From state 1 to 2, from 2 to 3, and all the rest until the final state, the forces were added between 13 pairs of cut sites on the linker staples (on the upper and lower edges), as illustrated by the red arrows in state 1. (b) Potential energy landscape associated with the applied spring forces for the cyclization. By extracting the initial and final potential energies (*i.e*., the potential energy of the springs) in a process from one state to the next, the external work is calculated. (c) Distance profile during the cyclization. This is the averaged distance between the cut sites of the linkers. The distance starts from ~84 nm (32 rows of dsDNA bundles or the height of the origami tile) and ends below 11 nm, indicating a cyclized conformation in the final state.

The total number of states were identified from the potential energy calculations. Figure 4b shows that each state has two (low and high) potential energies, except for states 0 and 10 (the initial and final states). The low energy is the ending potential energy of the calculation from the previous to the current state, while the high energy is the starting potential energy for the transition from the current state to the next. For example, the potential energies involved in the transition from state 2 to 3 are 7.5 kcal/mol and 5.86 kcal/mol, respectively. That is, the origami in state 2 was subject to a set of external forces with 7.5 kcal/mol, and the minimum potential energy of 5.86 kcal/mol was found after 2 million calculations. The difference (1.64 kcal/mol) is the mechanical energy into the DNA origami tile. If the forces were kept constant for further deformation, the upper/lower boundaries would not become closer. Therefore, a new stronger set of spring forces was needed (with higher force constant k and/or lower equilibrium length of the spring x0). For the transition from state 3 to 4, multiple potential energies were tried and 10 kcal/mol was found to further deform the origami without minimal excess energy. Calculations for other states were carried out similarly, by selecting the conformation at the potential minimum in the previous simulation as the starting point of the next computation. The simulations were halted when the upper/lower boundaries were close enough for the linkers to seal them together. Since each linker is 32 nt, the base-paired length is estimated as roughly 11 nm. Thus, 11 nm on average was used as the criterion for the distance between the boundaries. Overall, a total of 10 states were used in the simulations of cyclization.

Under spring forces, the origami tile gradually rolled up and deformed into a cylindrical shape. If a very high potential is given at once (e.g., 400 kcal/mol), the tile can cyclize immediately in a single computation (from the initial to the final states). However, such simulations will neither reveal the minimum energy needed for cyclization nor depict the details of the process. Instead, we used quasi-equilibrium deformations in the simulations. In the early states for cyclization (state 1 through 3), only the dsDNA bundles near the boundaries responded to the pulling forces (Figure 4a). The folding of the entire origami was not strong. In state 4, all the bundles start to twist and bend. This trend of deformation became more distinct in states 5 through 9. The global response to the forces suggests that the deformation in the origami propagates gradually. The structure in the final state did not change too much visually compared with that in state 9. The noticeable change is the alignment of dsDNA bundles in parallel positions. For all states from 1 to 9, the two boundaries are not parallel to each other, exhibiting a degree of the global twisting. Reaching the final state, they become aligned, indicating that the curvature is fully overcome and the tile cyclizes into a tube as anticipated.

In Figure 4b, the lines from one state to the next indicate the energy minimization process, while the vertical lines indicate the increase of the potential energy by introducing new sets of stronger spring forces. All the potential energy changes are added together, and the energy needed for the full cyclization is calculated as ~33.1 kcal/mol. To quantitatively illustrate the cyclization, the averaged distance between the cut site of the linkers on the upper and lower boundaries is plotted in Figure 4c. The distance starts at approximately 84 nm, which is consistent with the designed height (~88 nm). As the tile cyclizes, the distance reduces steadily and reaches around 10 nm in the final state. State 10 has ~63 nm in width and ~25 nm in diameter, which is comparable to the previous measurement of a cyclized tile (~65×45 nm for a flattened tube).^27^ This confirms the validity of our approach for cyclization in MD simulations.

### Energy Driven Planarization

In the cyclization, two deformation modes, overcoming the initial twist and flexing the tile into a tube, occur simultaneously. It is difficult to separate the two modes. Therefore, we perform MD simulations to study a deformation that purely overcomes the twisting (*i.e*., flattening). This will also verify the conceptual two-step process in our previous study.^27^ All of the 13,360 nucleotides were pulled to the imaginary center flat plane (dashed lines in Figure 5a) by applying the spring forces in a series of quasi-equilibrium deformations. Since the middle part of the origami tile is closer to flat compared with four corners, the imaginary center plane is placed through the middle. Then, strong forces are applied to the middle part in order to hold it in place, while the corners are subject to smaller flattening forces such that they will approach the center plane gradually and steadily. For example, the forces on dsDNA bundle number 12 through 21 are about 3 times stronger than those on the rest of the bundles, when transitioning from state 0 to state 1. If much stronger forces were applied at the corners, they would be compressed quickly and might be folded instead of moving toward the center plane. In states 2, 3 and 6, for example, the upper corners hovered on top of the flat part due to thermal fluctuation. They were allowed to deform toward the center plane due to small forces applied instead of folding locally. Moreover, much more energy would have been consumed, thus we could not determine the minimum energy for flattening. Figure 5a shows the gradual deformations of the tile from the twisted curvature to the flat conformation. We observe that the two upper corners along with the top few dsDNA bundles were not in the center plane until the last state. It is worth noting that a step to find the starting conformation was not needed in the flattening, because the origami tile does not have two directions to deform – it deforms only toward the center plane.

**Figure 5.**
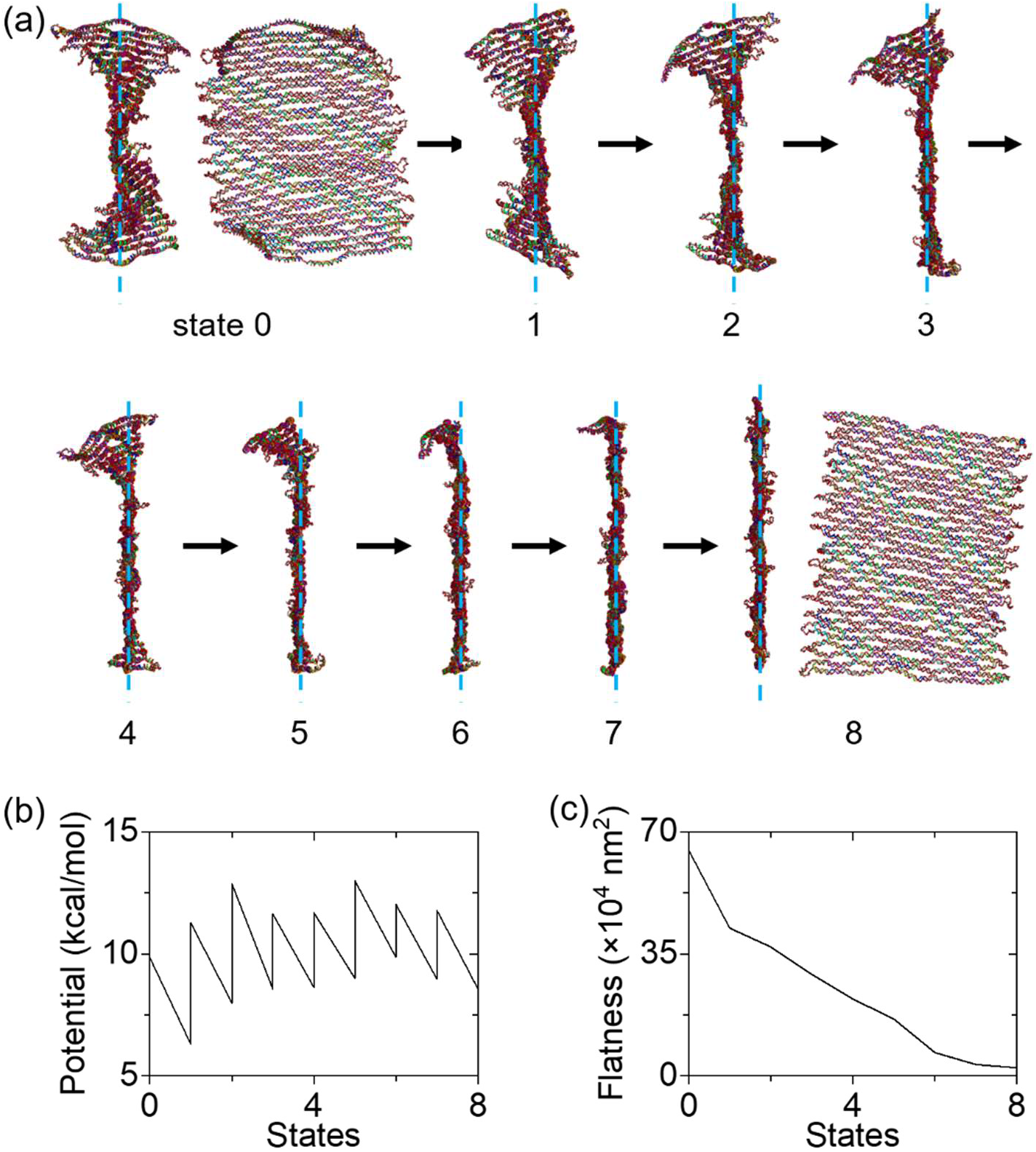
Flattening process of a DNA origami tile in MD simulations. (a) The origami tile flattens from the initial state (state 0) and undergoes gradual planarization until the final state (state 8). The flattening is induced by pulling all the nucleotides to the imaginary flat plane at the center (dashed turquoise line) with hypothetical springs. In the final state, the origami tile is planar. (b) External potential landscape during the flattening process. The external work was calculated in a similar approach as in the cyclization process. (c) Flatness during the flattening. This is a sum of the distance squared of each nucleotide to the imaginary flat plane. Given the radius of a dsDNA bundle (~1.1 nm), the flatness of a planar tile is ~2×10^4^ nm^2^. During the deformation, the flatness starts from ~6.5×10^5^ nm^2^ and ends at ~2.3×10^4^ nm^2^.

Similar to the cyclization process, the structure was deformed in a multistep process for flattening. The external forces and structural deformation were halted when the origami became flat enough. To develop a criterion, we defined the flatness as the sum of the distance squared of each nucleotide to the imaginary flat plane. For example, 0 flatness means perfectly flat. Since the origami tile has a non-zero thickness, its flatness is always greater than 0. The radius of regular, right-handed B-DNA is 1.1 nm. The flatness is ~2×10^4^ nm^2^ if we consider a reference state where all the nucleotides are 1.1 nm from the imaginary center plane. When the flatness of the origami tile is below 2.5×10^4^ nm^2^, the simulations were halted.

The rectangular origami tile gradually flattened with spring forces toward the plane. As shown in Figure 5a, all parts of the origami response to the forces in all the states. For all states from 0 to 8, the middle part stays in the center plane while all the corners gradually approach the plane. The energy is calculated in the same method as for the cyclization in Figure 4. The potential energy landscape and the flatness suggest that about 26.3 kcal/mol is required to flatten the origami tile (Figure 5b and c). During the flattening, the flatness starts from ~6.5×10^5^ nm^2^ and ends at ~2.3×10^4^ nm^2^ (within the criterion) The conformation in the final state measures around 86 nm in height and 63 nm in width, showing good agreement with our design and AFM imaging results.^27^

## DISCUSSION

### Equilibrium Conformations in CanDo and OxDNA

The results in Figure 3c and the initial state (state 0) in Figure 4a are slightly different. The dsDNA bundles are closely packed in the structure from CanDo, whereas gaps are seen in the oxDNA results. Thus, the origami height in Figure 3c appears to be shorter than that in Figure 4a. Several factors are accounted for these differences. FEM simulations in CanDo have the single base-pair resolution, treating ssDNA and crossovers as a connection without any length. These make the DNA origami stiffer than it is. For example, the crossovers are seen in Figure 3c since they have no length. Similarly, the unpaired scaffold domains at either side of the tile are not shown as fluffy segments. In contrast, oxDNA adopts the coarse-grained MD model, where each nucleotide is the basic group of atoms (*i.e*., single nucleotide resolution). The MD model accounts for several parameters in the simulations, including temperature, salt concentration, and major/minor groove of dsDNA. OxDNA provides significantly more detailed information than CanDo. For example, the neighboring dsDNA bundles connected by crossovers will stay together. However, the segments of neighboring bundles with no crossovers will show gaps in oxDNA, indicating the repulsion between dsDNA helices due to negative charges. This is not depicted in CanDo. Because of the gaps, the height in oxDNA is greater than that of CanDo, causing the difference in size. Aside from these differences, both conformations are comparable.

### Configuration of a Cyclized Origami Tile

The final configuration of the origami tile in Figure 4a slightly differs from that in Figure 3d. There are several reasons for the difference. The schematics in Figure 3d show the ideal configuration of the cyclized structure, where all the dsDNA bundles are perfectly aligned. In the oxDNA, however, the dsDNA bundles are bent slightly, the unpaired ss-scaffold segments cause attraction or repulsion forces, and the thermal fluctuation changes the overall conformation to some degree. The differences are limited, nevertheless. The cross-section of the MD simulation results resembles a circular tube as in Figure 3d. The upper and lower surface of the tube are parallel to each other. Finally, the size of the tube (about 63 nm in width and 25 nm in diameter) also agrees with the previous experimental results.^27^

### Deformation Process of Cyclization

Figure 4a shows one possible deformation pathway of cyclization with minimal energy. Overcoming of the twisting and bending of the structure occur simultaneously. The simulations also reveal the gradual propagation of the deformation in the structure during the cyclization. This information is available only with computational studies, whereas experiments show the origami conformations before and after cyclization. In addition, the theoretical approach cannot consider the cyclization process directly, rather it introduces a hypothetical intermediate step. Thus, MD simulations can provide detailed information such as the conformation and necessary energy during the cyclization process as well as pathways.

### Free Energy Change in Cyclization

The cyclization process was studied experimentally and theoretically in our previous report^27^ as well as by MD simulations in this work. As discussed above, the cyclization was theorized into two independent processes of flattening and flexing by introducing a hypothetical intermediate planar conformation. The energy for cyclization equals the summation of energy changes for flattening and flexing:

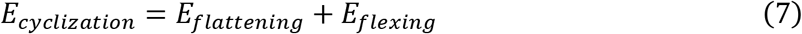

Figure 6 summarizes the cyclization and the conceptual two-step process. The energies extracted from all three approaches are in remarkable agreement. Note that it was impossible to determine the energies needed for flattening or flexing in experiments. The mechanical analysis and MD computation yielded *E_flattening_* of approximately 25.6 and 26.3 kcal/mol, respectively. It is worth noting that flexing was not performed in MD simulations, because the origami could twist again once subjected under flexing forces. For any given initial and final states, various deformation pathways may be proposed to calculate the related energies. However, whether the process is realistic or not may not be revealed by the theory alone, even if it is a free energy driven reaction. Therefore, performing simulations along with the theory will be beneficial.

**Figure 6.**
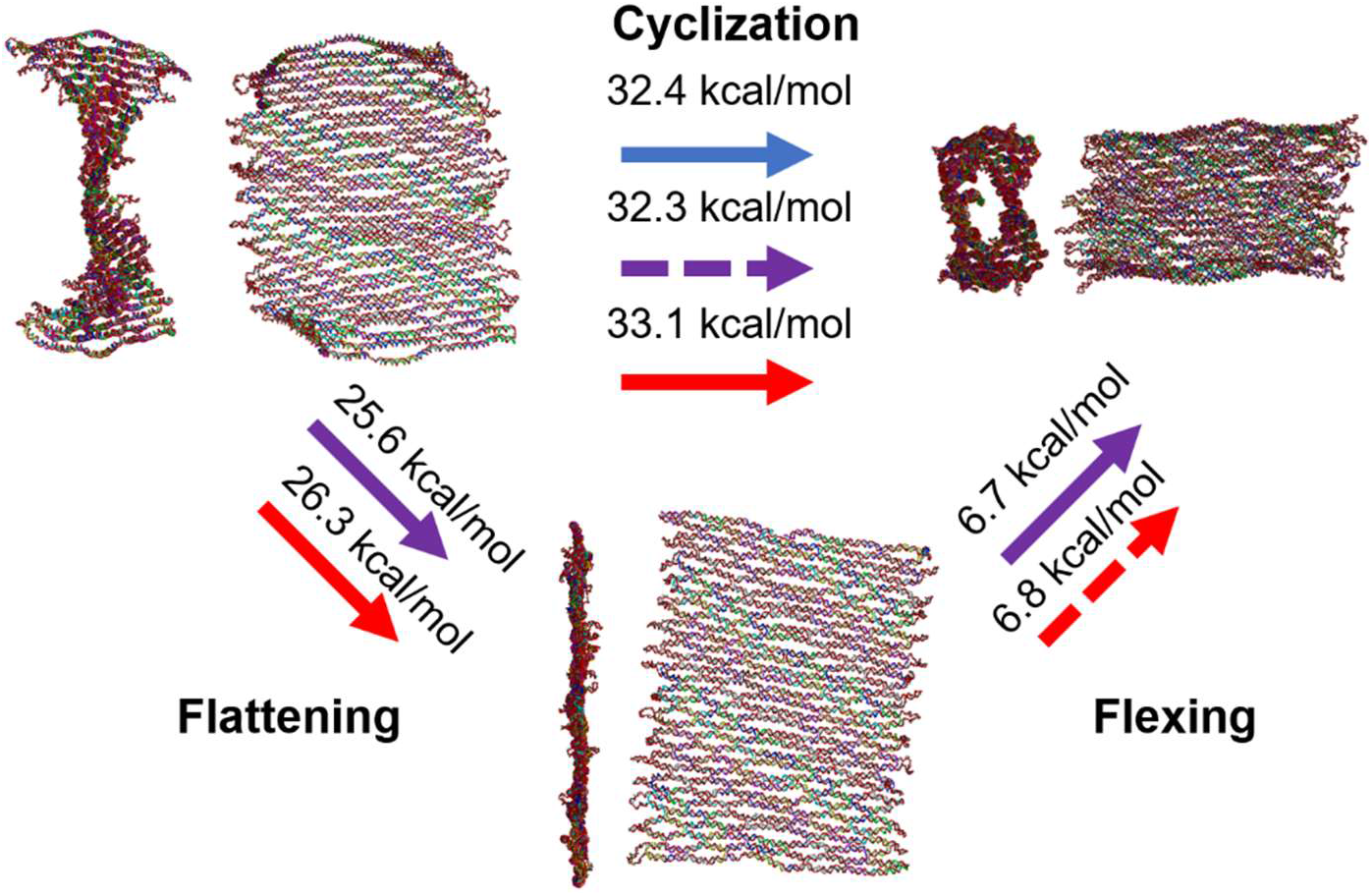
Overall analysis of the cyclization process. The cyclization of a DNA origami tile with a twist may be conceptualized as a two-step (flattening and flexing) process by introducing a hypothetical intermediate planar conformation. Blue, purple, and red arrows indicate the experiment, theory, and simulation, respectively. Solid arrows denote the direct results, while dashed arrows mark the indirect results from summation or subtraction of direct results. Experimentally determined free energy change for cyclization is approximately 32.4 kcal/mol. The theoretical value for the cyclization energy is obtained as ~32.3 kcal/mol by adding energies for flattening and flexing steps. The MD simulations yield a cyclization energy of about 33.1 kcal/mol, consistent with the experimental and theoretical values.

Since direct numerical studies on flexing are not practical, the flexing energy may be obtained from equation 1 using the cyclization and flattening energies, which is about 6.8 kcal/mol. This value also agrees well with the theoretical value of ~6.7 kcal/mol. The combined experiment, theory, and computation all agree well and verify the effectiveness of the simulation approach in this work.

In summary, coarse-grained MD simulations were performed on the oxDNA platform to provide critical insights on the cyclization of a DNA origami tile with initial twisting. This work advances our understanding of fundamental mechanics of DNA origami in general and should be useful for energy driven processes including adaptive reconfiguration, mechanical (or other) energy absorption mechanisms, and the interactions with other biomolecules.

## ACKNOWLEDGEMENT

This work was supported by the U.S. Department of Energy (DOE), Office of Science, Basic Energy Sciences (BES), under award no. DE-SC0020673 (computational studies) and by the U.S. National Science Foundation (NSF) under award no. 1710344 (experimental analysis).

## REFERENCES

1 Seeman, N. C. & Sleiman, H. F. DNA Nanotechnology. Nature Reviews Materials 3, 1–23 (2017).

2 Pinheiro, A. V., Han, D., Shih, W. M. & Yan, H. Challenges and Opportunities for Structural DNA Nanotechnology. Nature Nanotechnology 6, 763–772 (2011).

3 Seeman, N. C. Nucleic Acid Junctions and Lattices. Journal of Theoretical Biology 99, 237–247 (1982).

4 Lin, C., Liu, Y., Rinker, S. & Yan, H. DNA Tile Based Self - Assembly: Building Complex Nanoarchitectures. ChemPhysChem 7, 1641–1647 (2006).

5 Park, S. H., Pistol, C., Ahn, S. J., Reif, J. H., Lebeck, A. R., Dwyer, C. & LaBean, T. H. Finite-Size, Fully Addressable DNA Tile Lattices Formed by Hierarchical Assembly Procedures. Angewandte Chemie 118, 749–753 (2006).

6 Wei, B., Dai, M. & Yin, P. Complex Shapes Self-Assembled from Single-Stranded DNA Tiles. Nature 485, 623–626 (2012).

7 Ke, Y., Ong, L. L., Shih, W. M. & Yin, P. Three-Dimensional Structures Self-Assembled from DNA Bricks. science 338, 1177–1183 (2012).

8 Ke, Y., Ong, L. L., Sun, W., Song, J., Dong, M., Shih, W. M. & Yin, P. DNA Brick Crystals with Prescribed Depths. Nature Chemistry 6, 994–1002 (2014).

9 Yurke, B., Turberfield, A. J., Mills, A. P., Simmel, F. C. & Neumann, J. L. A DNA-Fuelled Molecular Machine Made of DNA. Nature 406, 605–608 (2000).

10 Liu, M., Fu, J., Hejesen, C., Yang, Y., Woodbury, N. W., Gothelf, K., Liu, Y. & Yan, H. A DNA Tweezer-Actuated Enzyme Nanoreactor. Nature Communications 4, 1–5 (2013).

11 Omabegho, T., Sha, R. & Seeman, N. C. A Bipedal DNA Brownian Motor with Coordinated Legs. Science 324, 67–71 (2009).

12 Lund, K., Manzo, A. J., Dabby, N., Michelotti, N., Johnson-Buck, A., Nangreave, J., Taylor, S., Pei, R., Stojanovic, M. N. & Walter, N. G. Molecular Robots Guided by Prescriptive Landscapes. Nature 465, 206–210 (2010).

13 Pan, J., Cha, T.-G., Li, F., Chen, H., Bragg, N. A. & Choi, J. H. Visible/Near-Infrared Subdiffraction Imaging Reveals the Stochastic Nature of DNA Walkers. Science Advances 3, e1601600 (2017).

14 Pan, J., Du, Y., Qiu, H., Upton, L. R., Li, F. & Choi, J. H. Mimicking Chemotactic Cell Migration with DNA Programmable Synthetic Vesicles. Nano Letters 19, 9138–9144 (2019).

15 Zhang, D. Y. & Seelig, G. Dynamic DNA Nanotechnology Using Strand-Displacement Reactions. Nature Chemistry 3, 103–113 (2011).

16 Li, F., Chen, H., Pan, J., Cha, T.-G., Medintz, I. L. & Choi, J. H. A DNAzyme-Mediated Logic Gate for Programming Molecular Capture and Release on DNA Origami. Chemical Communications 52, 8369–8372 (2016).

17 Zhang, D. Y. & Winfree, E. Control of DNA Strand Displacement Kinetics Using Toehold Exchange. Journal of the American Chemical Society 131, 17303–17314 (2009).

18 Rothemund, P. W. K. Folding DNA to Create Nanoscale Shapes and Patterns. Nature 440, 297–302 (2006).

19 Grima, J. N., Mizzi, L., Azzopardi, K. M. & Gatt, R. Auxetic Perforated Mechanical Metamaterials with Randomly Oriented Cuts. Advanced Materials 28, 385–389 (2016).

20 Castro, C. E., Kilchherr, F., Kim, D.-N., Shiao, E. L., Wauer, T., Wortmann, P., Bathe, M. & Dietz, H. A Primer to Scaffolded DNA Origami. Nature Methods 8, 221 (2011).

21 Kopperger, E., List, J., Madhira, S., Rothfischer, F., Lamb, D. C. & Simmel, F. C. A Self-Assembled Nanoscale Robotic Arm Controlled by Electric Fields. Science 359, 296–301 (2018).

22 Choi, J., Chen, H., Li, F., Yang, L., Kim, S. S., Naik, R. R., Ye, P. D. & Choi, J. H. Nanomanufacturing of 2D Transition Metal Dichalcogenide Materials Using Self - Assembled DNA Nanotubes. Small 11, 5520–5527 (2015).

23 Li, R., Chen, H. & Choi, J. H. Topological Assembly of a Deployable Hoberman Flight Ring from DNA. Small 17, 2007069 (2021).

24 Chen, H., Zhang, H., Pan, J., Cha, T.-G., Li, S., Andréasson, J. & Choi, J. H. Dynamic and Progressive Control of DNA Origami Conformation by Modulating DNA Helicity with Chemical Adducts. ACS Nano 10, 4989–4996 (2016).

25 Czogalla, A., Petrov, E. P., Kauert, D. J., Uzunova, V., Zhang, Y., Seidel, R. & Schwille, P. Switchable Domain Partitioning and Diffusion of DNA Origami Rods on Membranes. Faraday discussions 161, 31–43 (2013).

26 Mishra, S., Feng, Y., Endo, M. & Sugiyama, H. Advances in DNA Origami–Cell Interfaces. ChemBioChem 21, 33–44 (2020).

27 Chen, H., Weng, T.-W., Riccitelli, M. M., Cui, Y., Irudayaraj, J. & Choi, J. H. Understanding the Mechanical Properties of DNA Origami Tiles and Controlling the Kinetics of Their Folding and Unfolding Reconfiguration. Journal of the American Chemical Society 136, 6995–7005 (2014).

28 Li, Z., Liu, M., Wang, L., Nangreave, J., Yan, H. & Liu, Y. Molecular Behavior of DNA Origami in Higher-Order Self-Assembly. Journal of the American Chemical Society 132, 13545–13552 (2010).

29 Liber, M., Tomov, T. E., Tsukanov, R., Berger, Y. & Nir, E. A Bipedal DNA Motor that Travels Back and Forth Between Two DNA Origami Tiles. Small 11, 568–575 (2015).

30 Zhao, Z., Yan, H. & Liu, Y. A Route to Scale up DNA Origami Using DNA Tiles as Folding Staples. Angewandte Chemie 122, 1456–1459 (2010).

31 Andersen, E. S., Dong, M., Nielsen, M. M., Jahn, K., Subramani, R., Mamdouh, W., Golas, M. M., Sander, B., Stark, H. & Oliveira, C. L. Self-Assembly of a Nanoscale DNA Box with a Controllable Lid. Nature 459, 73–76 (2009).

32 Zadegan, R. M., Jepsen, M. D., Thomsen, K. E., Okholm, A. H., Schaffert, D. H., Andersen, E. S., Birkedal, V. & Kjems, J. Construction of a 4 Zeptoliters Switchable 3D DNA Box Origami. ACS Nano 6, 10050–10053 (2012).

33 Kuzuya, A. & Komiyama, M. Design and Construction of a Box-Shaped 3D-DNA Origami. Chemical Communications, 4182–4184 (2009).

34 Zhang, F., Jiang, S., Wu, S., Li, Y., Mao, C., Liu, Y. & Yan, H. Complex Wireframe DNA Origami Nanostructures with Multi-Arm Junction Vertices. Nature Nanotechnology 10, 779–784 (2015).

35 Benson, E., Mohammed, A., Rayneau-Kirkhope, D., Gådin, A., Orponen, P. & Högberg, B. Effects of Design Choices on the Stiffness of Wireframe DNA Origami Structures. ACS nano 12, 9291–9299 (2018).

36 Veneziano, R., Ratanalert, S., Zhang, K., Zhang, F., Yan, H., Chiu, W. & Bathe, M. Designer Nanoscale DNA Assemblies Programmed from the Top Down. Science 352, 1534–1534 (2016).

37 Ke, Y., Castro, C. & Choi, J. H. Structural DNA Nanotechnology: Artificial Nanostructures for Biomedical Research. Annual Review of Biomedical Engineering 20, 375–401 (2018).

38 Ijäs, H., Nummelin, S., Shen, B., Kostiainen, M. A. & Linko, V. Dynamic DNA Origami Devices: from Strand-Displacement Reactions to External-Stimuli Responsive Systems. International Journal of Molecular Sciences 19, 2114 (2018).

39 Zhang, Y., Pan, V., Li, X., Yang, X., Li, H., Wang, P. & Ke, Y. Dynamic DNA Structures. Small 15, 1900228 (2019).

40 Chen, H., Cha, T.-G., Pan, J. & Choi, J. H. Hierarchically Assembled DNA Origami Tubules with Reconfigurable Chirality. Nanotechnology 24, 435601 (2013).

41 Zhang, Z., Yang, Y., Pincet, F., Llaguno, M. C. & Lin, C. Placing and Shaping Liposomes with Reconfigurable DNA Nanocages. Nature Chemistry 9, 653–659 (2017).

42 Zhang, F., Nangreave, J., Liu, Y. & Yan, H. Reconfigurable DNA Origami to Generate Quasifractal Patterns. Nano letters 12, 3290–3295 (2012).

43 Kauert, D. J., Kurth, T., Liedl, T. & Seidel, R. Direct Mechanical Measurements Reveal the Material Properties of Three-Dimensional DNA Origami. Nano Letters 11, 5558–5563 (2011).

44 Oberstrass, F. C., Fernandes, L. E. & Bryant, Z. Torque Measurements Reveal Sequence-Specific Cooperative Transitions in Supercoiled DNA. Proceedings of the National Academy of Sciences 109, 6106–6111 (2012).

45 Sa-Ardyen, P., Vologodskii, A. V. & Seeman, N. C. The Flexibility of DNA Double Crossover Molecules. Biophysical Journal 84, 3829–3837 (2003).

46 Li, R., Chen, H. & Choi, J. H. Auxetic Two-Dimensional Nanostructures from DNA. Angewandte Chemie, doi:10.1002/anie.202014729 (2021).

47 Goodman, R. P., Schaap, I. A., Tardin, C. F., Erben, C. M., Berry, R. M., Schmidt, C. F. & Turberfield, A. J. Rapid Chiral Assembly of Rigid DNA Building Blocks for Molecular Nanofabrication. Science 310, 1661–1665 (2005).

48 Douglas, S. M., Marblestone, A. H., Teerapittayanon, S., Vazquez, A., Church, G. M. & Shih, W. M. Rapid Prototyping of 3D DNA-Origami Shapes with CaDNAno. Nucleic Acids Research 37, 5001–5006 (2009).

49 Dietz, H., Douglas, S. M. & Shih, W. M. Folding DNA into Twisted and Curved Nanoscale Shapes. Science 325, 725–730 (2009).

50 Chen, H., Li, R., Li, S., Andréasson, J. & Choi, J. H. Conformational Effects of UV Light on DNA Origami. Journal of the American Chemical Society 139, 1380–1383 (2017).

51 Ke, Y., Bellot, G., Voigt, N. V., Fradkov, E. & Shih, W. M. Two Design Strategies for Enhancement of Multilayer–DNA-Origami Folding: Underwinding for Specific Intercalator Rescue and Staple-Break Positioning. Chemical Science 3, 2587–2597 (2012).

52 Zadegan, R. M., Lindau, E. G., Klein, W. P., Green, C., Graugnard, E., Yurke, B., Kuang, W. & Hughes, W. L. Twisting of DNA Origami from Intercalators. Scientific Reports 7, 1–5 (2017).

53 Snodin, B. E. K., Randisi, F., Mosayebi, M., Šulc, P., Schreck, J. S., Romano, F., Ouldridge, T. E., Tsukanov, R., Nir, E., Louis, A. A. & Doye, J. P. K. Introducing Improved Structural Properties and Salt Dependence into a Coarse-Grained Model of DNA. Journal of Chemical Physics 142, 234901 (2015).

54 Engel, M. C., Smith, D. M., Jobst, M. A., Sajfutdinow, M., Liedl, T., Romano, F., Rovigatti, L., Louis, A. A. & Doye, J. P. K. Force-Induced Unravelling of DNA Origami. ACS Nano 12, 6734–6747 (2018).

