## Supporting Information for "Elucidating the Mechanical Energy for Cyclization of a DNA Origami Tile"

**Content**

S1. Starting Conformation for Cyclization
S2. Potential Energy Landscape

**S1. Starting Conformation for Cyclization**

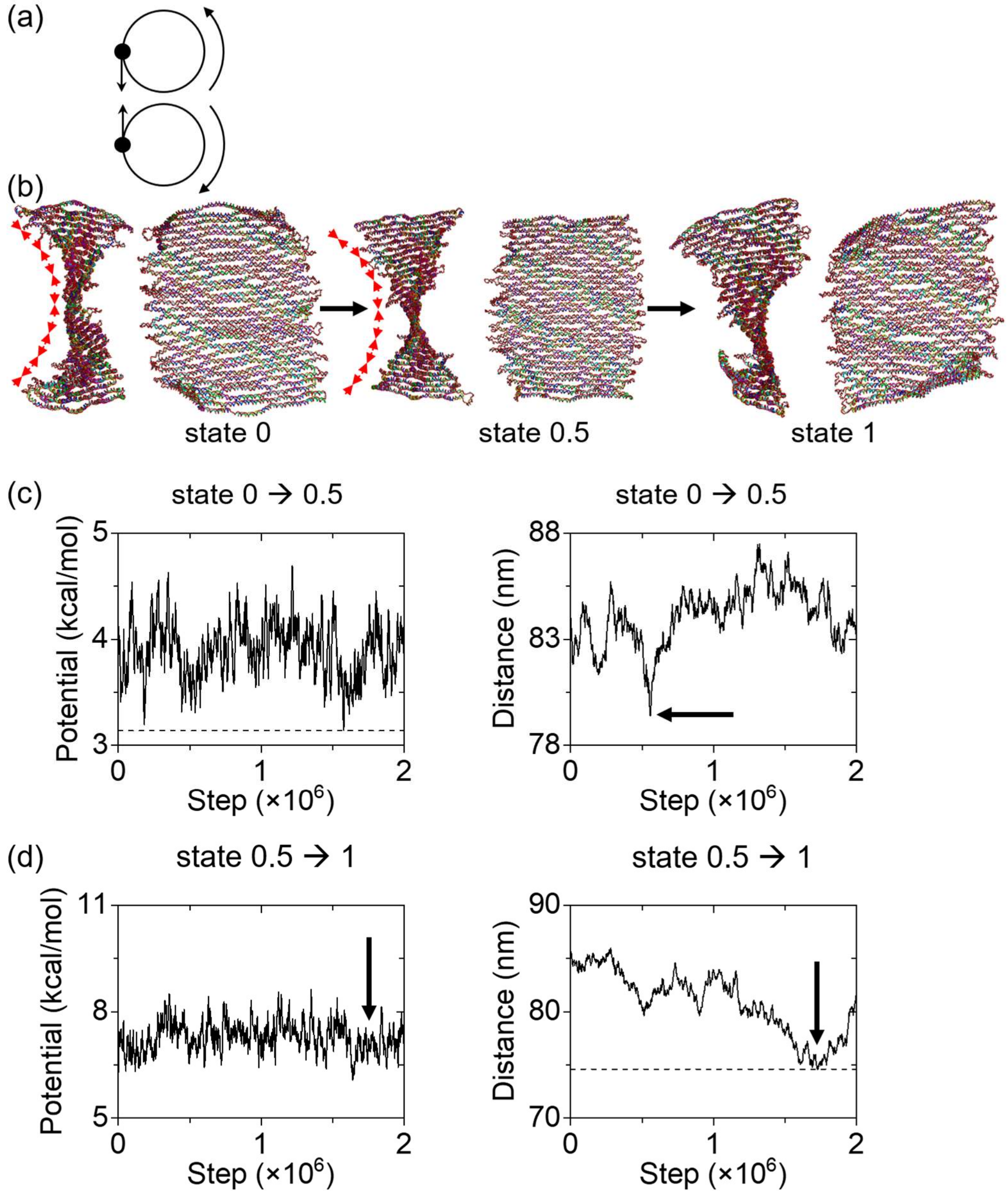


**Figure S1.** (a) Loading of hypothetical linear spring forces (arrows on the left side) in the side view of two neighboring dsDNA bundles in the single-layer DNA origami rectangle. Solid dots indicate the nucleotides on two adjacent bundles that are located at the very left side of the bundles. By pulling them toward each other, the bundles will rotate in the directions noted by the arrows on the right side. Thus, the structure will only deform toward the left side when subjected to the forces on 13 pairs of cut side of the linkers. (b) Finding the starting conformation of the DNA origami tile for cyclization. The DNA origami tile is under the applied forces between all the neighboring dsDNA bundles. States 0 and 1 are the same as in Figure 4. Upon the external forces for cyclization, the structure may fold in a randomly direction on its either side. To ensure the cyclization in a particular direction, we added an intermediate state (termed, state 0.5) between state 0 and 1. Thus, we have two processes of deformation to reach the starting conformation (state 1). Red arrows in states 0 and 0.5 show the forces, which pulls the bundles together only on the left side of the origami. (c) Potential energy landscape associated with the applied forces and the distance between the upper and lower boundaries from state 0 to 0.5. The structure that fold only on the left side will have the minimum distance between upper and lower boundaries. The minimum distance from state 0 to 0.5 is found at step number 0.558 million (indicated by the black arrow). However, this conformation did not result in cyclized deformations under subsequent loading on the 13 linkers. Instead, we chose the minimum potential at step number 1.58 million and the conformation at this step was used for state 0.5. The energy into the origami tile from state 0 to 0.5 is 0.82 kcal/mol. (d) Potential energy landscape and the distance between the upper and lower edges from state 0.5 to 1. The minimum distance was reached at step number 1.732 million. We found that the structure led to cyclized deformations under subsequent loading on the 13 linkers. Thus, the conformation at this step was chosen as state 1. In this process, the mechanical energy applied on the origami tile is -1.08 kcal/mol (the potential indicated by the black arrow).

In the equilibrium, the DNA origami tile has a twisted curvature as shown in Figure S1a. Applying spring forces on the linkers for tile cyclization directly from state 0 may cause the this single-layer DNA origami tile to fold on either side. That is, simply applying the forces may not necessarily result in cyclization in a particular direction (*e.g.*, folded towards the left), and one may find that part of the origami structure cyclizes towards the left and other parts of the tile folds towards the right. This is because the origami tile can deform and fold on its either side with no preference. To resolve this issue, we pulled the neighboring bundles on one side of the origami such that the tile will deform and cycle only in that direction upon the subsequent loading on the 13 linkers (*e.g.*, on the left side as illustrated in Figure S1a). The structure that folds only on one side is named 'starting conformation' in state 1. Note that states 0 and 1 in Figure S1a are the same as in Figure 4.

The hypothetical linear spring forces were applied to selected sites on all the neighboring DNA bundles to achieve the starting conformation. The selected sites are the nucleotides at the very left side of the origami tile as shown in Figure S1a. In total, there are 1113 pairs of loading. Since the structure that leads to cyclized deformations toward the left side was not found by a single process from state 0 to 1, we added an intermediate state (state 0.5) between state 0 and 1. Thus, we have two processes of deformation to reach the starting conformation. The structure that fold only on the left side should have the minimum distance between upper and lower boundaries. However, we found that the minimum distance from state 0 to 0.5 (black arrow) did not result in cyclized deformations under subsequent loading on the linkers. Therefore, state 0.5 was selected as the conformation that yields the minimum potential energy (step number 1.58 million) rather than the minimum distance (step number 0.558 million). By applying stronger forces on 1113 pairs (*i.e.*, larger spring constant *k*), state 0.5 was transformed to state 1. The minimum distance from state 0.5 to 1 was found at step number 1.732 million, and this conformation was found to deform toward a cyclized tube under loading on the 13 linkers. The total energy consumed from state 0 to 1 is approximately -0.26 kcal/mol. The negative sign implies that the energy was gained from the system during the deformation. This energy gain is due to thermal fluctuation, since the only source of applied energy to the system is the spring forces.

**S2. Potential Energy Landscape**

Quasi-equilibrium deformations were simulated in oxDNA to find the minimum of global potential energy in a series of states. In the transition from a state to the next, the calculation step number that resulted in the minimum potential energy. The DNA origami conformation at that step was then used for the next state calculations. In Figure S2, we present the potential energy landscape of the cyclization process and how the global minimum was selected.

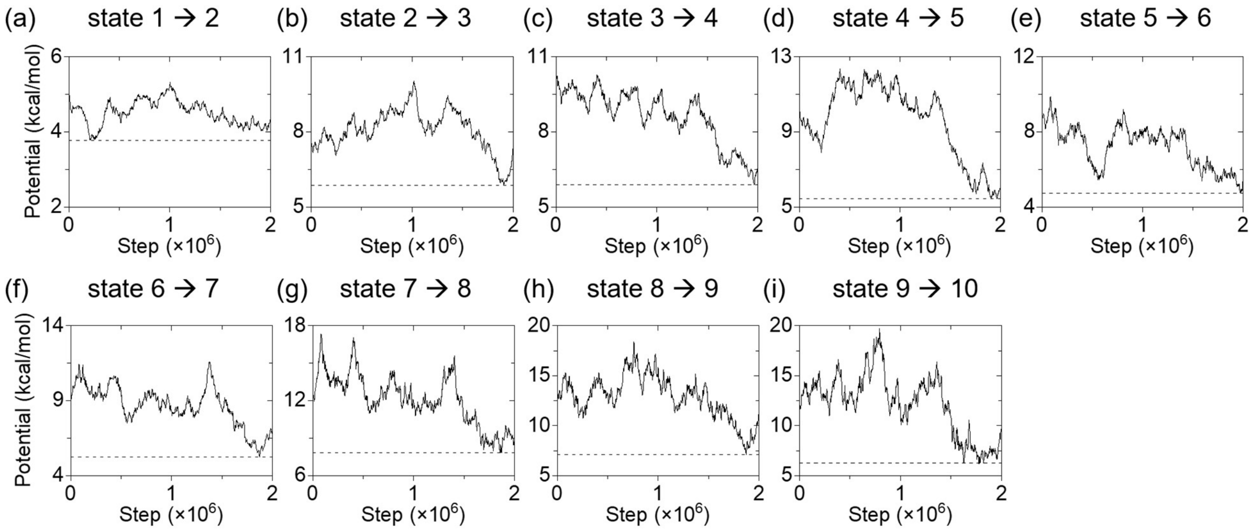


**Figure S2.** Potential energy landscape for the cyclization of the DNA origami tile. The origami tile was subject to spring forces for 2 million steps from a state to the next. The solid lines represent the potential energies from MD computation, while the dashed lines indicate the global minimum. The point where the solid line meets the dashed line indicates the position of the selected conformation for the next state.

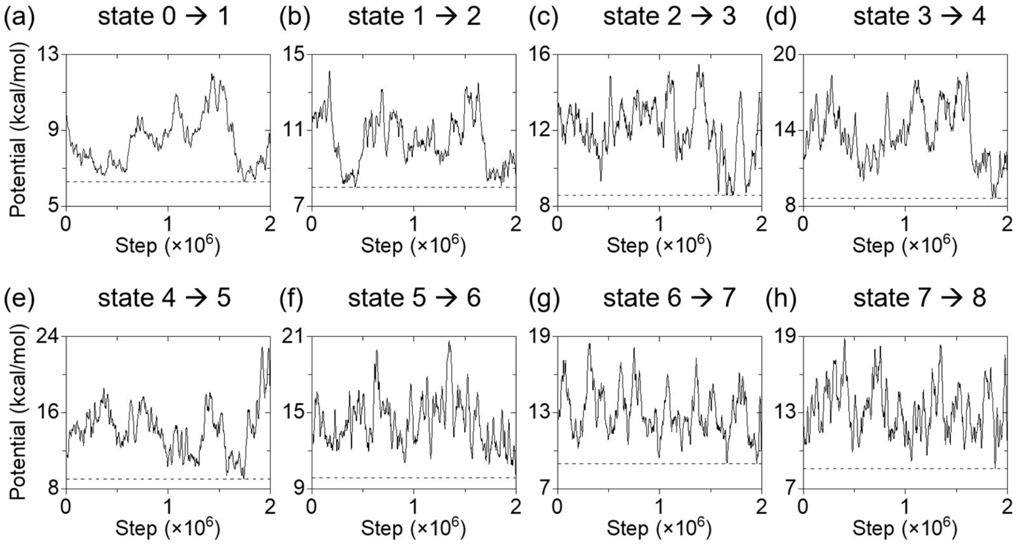


**Figure S3.** Potential energy landscape for the flattening of the DNA origami tile. The notation and the selection of the conformation for the next state are the same as Figure S2.
